# The role of sodium thiocyanate supplementation during dextran sodium sulphate-stimulated experimental colitis

**DOI:** 10.1101/2020.05.27.117994

**Authors:** Yuyang (Anna) Liu, Thomas Burton, Benjamin Saul Rayner, Patrick San Gabriel, Han Shi, Mary El Kazzi, XiaoSuo Wang, Joanne M Dennis, Gulfam Ahmad, Paul Kenneth Witting, Belal Chami

## Abstract

Ulcerative colitis is a condition characterised by the infiltration of leukocytes into the gastrointestinal wall. Leukocyte-MPO catalyses hypochlorous acid (HOCl) and hypothiocyanous acid (HOSCN) formation from chloride (Cl^-^) and thiocyanous (SCN^-^) anions, respectively. While HOCl indiscriminately oxidises biomolecules, HOSCN primarily targets low-molecular weight protein thiols. Oxidative damage mediated by HOSCN may be reversible, potentially decreasing MPO-associated host tissue destruction. This study investigated the effect of SCN^-^ supplementation in a model of acute colitis. Female mice were supplemented dextran sodium sulphate (DSS, 3% w/v) in the presence of 10 mM Cl^-^ or SCN^-^ in drinking water *ad libitum*, or with salts (NaCl and NaSCN only) or water only (controls). Behavioural studies showed mice tolerated NaSCN and NaCl-treated water with water-seeking frequency. Ion-exchange chromatography showed increased fecal and plasma SCN^-^ levels in thiocyanate supplemented mice; plasma SCN^-^ reached similar fold-increase for smokers. Overall there was no difference in weight loss and clinical score, mucin levels, crypt integrity and extent of cellular infiltration between DSS/SCN^-^ and DSS/Cl^-^ groups. Neutrophil recruitment remained unchanged in DSS-treated mice, as assessed by fecal calprotectin levels. Total thiol and tyrosine phosphatase activity remained unchanged between DSS/Cl^-^ and DSS/SCN^-^ groups, however, colonic tissue showed a trend in decreased 3-chlorotyrosine (1.5-fold reduction, *p*<0.051) and marked increase in colonic GCLC, the rate-limiting enzyme in glutathione synthesis. These data suggest that SCN^-^ administration can modulate MPO activity towards a HOSCN-specific pathway, however, this does not alter the development of colitis within a DSS murine model.

**Highlights:**

1. Sodium thiocyanate (SCN^-^) supplementation increased plasma and fecal SCN^-^ level.
2. Thiocyanate supplementation diverted HOCl-mediated oxidative damage in mice colon.
3. Thiocyanate supplementation stimulated thiol synthesis in the DSS colitis model.
4. Thiocyanate provides no protection in an acute experimental model of UC.

## 1. Introduction

Ulcerative colitis (UC) is a major pathological presentation of inflammatory bowel disease (IBD), a multifactorial immune response of the gastrointestinal (GI) tract, characterised by chronic leukocyte infiltration and remitting intestinal inflammation [1, 2]. Irreversible alterations in bowel structure and function are a hallmark feature of IBD, where it is increasingly recognised that reactive oxygen species (ROS) play a significant role in mediating host-tissue injury in the affected gastrointestinal tract.

Increasing evidence has identified the importance of ROS-mediated oxidative stress during inflammatory conditions such as UC [1, 3]. UC patients characteristically experience an exacerbated recruitment of polymorphonuclear neutrophils (PMNs) and macrophages into the colonic submucosa beneath active sites of ulceration [4]. Colonic and fecal myeloperoxidase (MPO), an enzyme produced and contained within PMNs, are significantly increased and correlate with disease severity in UC patients [5, 6], while others have shown an increase in colonic interleukin 8 (IL-8), a known activator of neutrophil MPO degranulation [6, 7].

The current view is that MPO is a major source of potent two-electron oxidants that include the hypohalous acids HOCl, HOSCN and hypobromous (HOBr), each derived from the corresponding (pseudo)halide substrates Cl^-^, SCN^-^ and Br^-^, respectively [6, 8]. HOCl is highly reactive and non-specifically targets a vast range of biomolecules including amine, amides, thiols, lipids, nucleic acids and numerous amino acids [9] with much of this HOCl-mediated damage is considered irreversible [9]. For example, 3-chlorotyrosine (3-Cl-Tyr), a specific oxidative marker of HOCl, is significantly increased in colonic and serum samples of patients with IBD [10]. In contrast, HOSCN preferentially oxides thiol groups in molecules such as glutathione (GSH) and GAPDH [11]. HOSCN-mediated thiol alteration is largely reversible as some of the oxidation products can be reduced by biological reductants to reform the parent thiol, thereby protecting thiols from permanent damage and in some cases restore enzymatic function [11–14].

Normal halide/pseudo-halide ion concentration in healthy human plasma ranges between 100-140 mM (Cl^-^); ~50 μM (SCN^-^) and 20-100 μM (Br^-^) [8]. Under normal physiological conditions, HOCl is the major ROS product of the MPO halogenation cycle owing to the relatively high concentration of Cl^-^ in biological systems [8]. In contrast, SCN^-^ has a much higher specificity for MPO than Cl^-^ (specificity constant; SCN:Cl:Br; 730:1:60 – a relative rate of substrate reaction with compound I) [15]. However, the relatively low plasma SCN^-^ concentration, compared to that of Cl^-^ renders lower HOSCN production from the halogenating cycle. Nevertheless, SCN^-^ has the potential to become a competitive substrate for MPO when its level is elevated beyond normal plasma concentrations. In addition to its 730-fold higher substrate specificity for MPO, SCN^-^ also exhibits a higher reaction rate than other halides, with the rate constants of reaction for SCN^-^, Cl^-^ and Br^-^ with MPO measured as 9.6 × 10^6^; 2.5 × 10^4^ and 1.1 × 10^6^ M^-□1^ s^-□1^, respectively [9, 15, 16].

SCN supplementation has been a recently growing area of research to mitigate the effects of MPO-derived HOCl in chronic inflammatory disease, based on population studies showing dietary and environmental SCN is protective in various pathologies [17]. For example, plasma SCN^-^ is associated with an all-cause decrease in human myocardial infarction [18]. Furthermore, increased bioavailability of SCN^-^ in airway epithelial cells in Cystic Fibrosis patients undergoing nebulized hypertonic saline therapy is thought to contribute increased biocidal activity [19].

Interestingly, active smokers show reduced risk and are protected from the clinical symptoms of UC [20–23]. Circulating SCN^-^ concentrations are significantly elevated in smokers compared to non-smokers, and plasma SCN^-^ values can exceed 200 μM (representing an approximate 2-fold increase in circulating SCN^-^), albeit with a mean value of 180 ± 55 μM [14, 24]. Thus, the goal of the study was to investigate if dietary SCN^-^ supplementation confers protective effects against experimental colitis through redirection of the MPO halogenation cycle to shift toward favoring HOSCN production above HOCl.

## 2. Materials and Methods

### 2.1 Chemicals

All laboratory chemicals were sourced from Sigma-Aldrich (Castle Hill, NSW, Australia and St Louis MO, USA) unless otherwise specified and were of the highest quality available at time of purchase. Protein analyses were performed using a commercial bicinchoninic acid assay [Sigma-Aldrich, Sydney Australia], and quantitative determinations were conducted with a standard curve prepared with bovine serum albumin as protein under identical experimental conditions.

### 2.2 IntelliCage experiments

All animal work was conducted at the animal house facility in the Charles Perkins Centre, The University of Sydney. To control the administration of variously treated drinking water to group-housed mice and to assess the functional effects of the treatment on general activity and water-seeking behaviour, we utilised the IntelliCage system (TSE Systems GmbH, Germany). Features of this system have been described elsewhere in detail [25, 26]. Briefly, animals were group-housed in an automated home-cage monitoring system where food was available *ad libitum* and access to water for each individual was controlled by the experimenter. Behavioural measures included 1) the number of visits to the four operant corners; 2) the number and duration of nosepokes (NPs) to the left or right ports within each operant corner (8 NP ports per IntelliCage) and; 3) the number of licks at the two water bottles in each operant corner (8 bottles per IntelliCage).

Prior to entering the IntelliCage, mice were implanted subcutaneously with a radio-frequency identification transponder (Datamars SA, Switzerland) under light isoflurane anaesthesia. Each IntelliCage housed 12 female mice for the duration of the experiment, with an even representation of the four treatment groups within each IntelliCage (i.e. n = 3 mice per treatment group per IntelliCage, pseudorandomly determined). Once animals were introduced into the IntelliCage, the behavioural measures defined above were quantified continuously for the duration of the experiment. The experiment proceeded as follows:

#### 2.2.1 Acclimatisation and Shaping

##### Free Adaptation (FA), 7 days

*Ad libitum* access to all 8 water bottles in the IntelliCage. Animals were free to visit all corners, NP at all NP ports and drink from each bottle.

##### Location Specific Nosepoke Adaptation (LS-NPA), 7 days

Mechanical doors at all NP ports barred access to the water bottles. A NP (i.e. snout breaking the IR beam of the port) could open the corresponding door for 8 seconds, allowing the animal to consume fluid from the water bottle. Only one door opening per visit to an operant corner was allowed. Importantly, in each IntelliCage only one bottle was accessible in a given corner for the remainder of the experiment. Hence, out of the eight possible reward sites only 4 were active. Furthermore, each individual animal could only access at one of the NP ports/water bottles. Therefore, 3 animals were allocated to each active reward site. Although drinking was restricted to one reward site for each individual for the remainder of the experiment, animals were free to visit and NP at all locations within the IntelliCage.

##### Salt Acclimatisation, 6 days

Same as LS-NPA except that 3 out of the 4 active water bottles per cage were filled with NaCl solution (NaCl dissolved in tap water) in increasing concentrations (Days 1-2: 2mM NaCl; Days 3-4: 6mM NaCl; Days 5-6: 10mM NaCl). Note that one bottle in each IntelliCage continued to contain tap water only (since the “control” group would receive tap water in the subsequent phase and would therefore not require salt acclimation). This was randomly determined.

### 2.3 DSS-induced acute colitis model

All animal work was approved by the Animal Ethics Committee, University of Sydney (AEC 2017/1119) and complied with the *New South Wales Animal Research Act 1985*. Female C57BL/6 wildtype mice aged 6 weeks old (n=24) [Animal Resource Centre, Canning Vale, Western Australia] were housed in two programmable automated IntelliCage (TSE Systems GmbH, Germany; (2 x 12 mice per cage) and individually tagged with an RFID transponder implanted subcutaneously. Mice were then trained to visit and nose poke to obtain water located at pseudorandomly determined corner of the IntelliCage over the course of 2-3 weeks (animals retained their previously allocated reward site throughout the experiment; see Supplementary section). The number and duration of corner visits, nose-pokes and licks for individual mice allocated to specific experimental groups were automatically recorded and this water-seeking behaviour quantified. Female mice were selected for experiments as they can be more readily cohoused in large groups than male mice who are prone to aggressive behaviour, particularly when encountering unfamiliar animals[27].

The drinking water of each mouse group (n=6) was treated with either water alone (control), 0.7 mM NaCl + 10 mM NaSCN (salts control), 3% w/v DSS + 10 mM NaCl (DSS/Cl^-^ group) and or 3% w/v DSS + 10 mM NaSCN (DSS/SCN^-^ group) over 8 days. Dextran sodium sulphate (DSS) [MP biomedicals, Seven Hills, Australia], sodium thiocyanate (NaSCN group) and sodium chloride (NaCl) were provided to mice in drinking water *ad libitum* and changed daily. Notably, Na^+^ ions present in supplemented DSS and NaSCN contributed to a salty taste in the drinking water. Therefore to account for this saltiness, research grade NaCl was added to generate the salts control and DSS/Cl^+^ groups in order to normalise the salt palatability of the drinking water. The concentration of NaCl added to drinking water was proportional to the concentration of Na^+^ in DSS and in NaSCN, respectively. Fecal samples were obtained on day 2 of DSS challenge. Animals were sacrificed via cervical dislocation following isoflurane induced anaesthesia. Upon sacrifice, blood was collected via cardiac puncture in EDTA-containing vials and plasma immediately prepared by centrifugation (13,000 rpm x *g*, 10 min, 4 °C), entire colons and fecal pellets and samples were stored at −80 °C until required for analysis. Where required, colon and fecal pellets were cut into small sections and homogenised in 1 mL lysis buffer with 50 mM phosphate (pH 7.4) containing: 1 mM EDTA, 10 mM butylated hydroxy toluene, 1 tablet/50 mL protease inhibitor cocktail (Roche, Basel, Switzerland), using a rotating piston and matching Teflon-covered tube (Wheaton, Millville, NJ) as described in detail previously [28, 29].

### 2.4 Ion-Exchange Chromatographic detection of SCN^-^ anions

Ion-exchange chromatography was performed on isolated plasma and homogenised fecal samples using the Dionex ICS-2100 System [Thermo Fisher Scientific, Australia] with an Anion Self-Regenerating Suppressor^®^ 300 operating at 124 mA, 35 °C. 12.5 μL injection was made from each sample; an IonPac™ AS 16 IC column and an AG 16 guard were then used to elute peaks with 50 mM KOH at 0.75 mL/min for 12 min [8, 14]. Ion peak areas were calculated using Cholmeron™ chromatography data system software (V. 7.1). Plasma and fecal SCN^-^ concentrations were determined based on a NaSCN standard curve (generated under identical conditions and using authentic NaSCN over a dose range 0-200 μM). Total SCN^-^ determined in fecal samples were normalised against corresponding total fecal protein levels as determined using a bicinchoninic assay.

### 2.5 Biochemical luminol-based myeloperoxidase assay

To determine the *ex vivo* inhibitory effect of SCN on MPO activity, the luminol biochemical assay was employed as previously described [30]. Briefly, a master mix containing the working concentration of 0.2μg/mL MPO and 0.8mM (w/v) luminol was supplemented with 67mM (w/v) NaCl. Varying concentrations of NaSCN were prepared and added to the master mix solution to a final concentration of 0.01, 0.1, 1 & 5 mM with subsequent agitation on an orbital shaker at 150 r.p.m for 20 min in a black flat-bottom 96-well plate. Following this, 45mM (v/v) H_2_O_2_ was added simultaneously to each well and the luminescence signal was immediately measured using a plater reader [Tecan, Austria (30063849)].

### 2.6 Liquid Chromatography-Mass Spectrometry assessment of 3-Cl-Tyrosine/Tyrosine ratios

Colonic levels of the HOCl biomarker 3-Cl-Tyr (expressed as the 3-Cl-Tyr/Tyr ratio) were measured based on an established protocol [31, 32]. Precipitation of colonic proteins from samples of colon homogenate (200 μL) were performed with 0.3 % w/v deoxycholic acid, 50% w/v trichloroacetic acid (TCA), 5% w/v TCA (100 μL) and icecold 100% acetone. Following addition of 4 M methanesulfonic acid containing 0.2% w/v tyrosine and internal standard mixture, samples were placed inside degassed hydrolysis reaction vessels and heated at 110 °C for 16 h for complete acid forming part hydrolysis. Solid phase extraction (SPE) was then performed using chromatography SPE cartridges [Supelclean Envi-Chrom, 250mg, St. Louis, USA] with 80% v/v methanol/H_2_O as eluent. The resulting eluate (100% methanol) was dried under vacuum and immediately reconstituted with 0.1% w/v formic acid in water (HPLC grade). Quantitative LC/MS was performed using a liquid chromatography triple quadrupole mass spectrometer 8050 [Shimadzu Corporation, Kyoto, Japan]. Components of each sample were separated using a LC Zorbax SB-Phenyl column [3.0 x 50 mm, 1.8 μm, Agilent Technologies, Santa Clara, USA and binary gradient elution method. Peaks were eluted over 15 min at a flow rate of 0.2 mL/min. Resultant chromatographic peaks were then evaluated and quantified using the LabSolution© software (Version 5.91,Shimadzu Corp,. Japan,). Labelled Tyrosine (L-Tyrosine-^13^C_9_,^15^N) and labelled 3-Cl-Tyrosine (L-3-Cl-Tyrosine-^13^C_9_,^15^N) were used as internal standards in both calibration curve and samples to quantify levels of native 3-Cl-Tyr and unmodified Tyr based on the corresponding standard curves generated on the same day of analysis under the same chromatographic conditions.

### 2.7 Enzyme-linked Immunosorbent Assay measurement of colon calprotectin

Calprotectin levels in clarified colon homogenates diluted to 10 μg total protein were measured using a sandwich enzyme-linked immunosorbent assay (ELISA) Kit [Hycult Biotech, USA (HK214-01)], as per manufacturer’s instructions. Colormetric development of 3,3’,5,5’-Tetramethylbenzidine was immediately read at 450 nm with a plater reader [Tecan, Austria (30063849)]) and the concentration determined by comparison with a standard curve prepared using authentic calprotectin provided in the commercial kit. All data were normalised to the corresponding protein level in the sample as expressed as μg/ mL homogenate of protein.

### 2.8 Plasma protein tyrosine phosphatase activity assay

Samples of plasma (5 μL) were diluted to 50 μL in a clear, flat-bottomed 96-well plate with 0.5 mM MgCl_2_ solution in PBS. Shrimp alkaline phosphatase (1:200 v/v) (Quantum Scientific) and nanopure water were used as positive and negative controls, respectively. Next, liquid para-nitro-phenyl phosphate (pNPP, 50 μL, dissolved in 50 mM phosphate buffer; pH 7.4, final concentration 25 μM) was added to all wells and the plate was incubated at 37 °C with absorbance measured in a plate reader (Tecan, Austria (30063849)] every 10 min at 405 nm for 20 h. The change in absorbance was plotted against time and a slope generated by linear regression (performed with Prism *V7*) over the first 3 h, which reflected plasma PTP activity and was normalised to the corresponding protein level in the sample, expressed as units/mg homogenate protein.

### 2.9 DTNB thiol Assay

Plasma thiol levels were detected via reaction with 5,5’-dithiobis(2-nitrobenzoic acid) (DTNB) [31].5 μL of plasma samples were loaded in triplicates and made up to 10 μL with nanopure water. A series of standards ranging from 0 mM to 0.5 mM were also prepared from a 0.5 mM glutathione (GSH) stock solution [Sigma Aldrich, USA (G4251)]. Phosphate buffer (100 mM) was made from 16 mL of 6.8 g of KH_2_PO_4_ [BHD, 10203] in 500 mL of nanopure and 84 mL of 11.4 g of K_2_HPO_4_ [Sigma Aldrich, USA (P-8281)] in 500 mL of nanopure water and adjusted to pH 7.4. Standards and duplicates of each sample were loaded with 200 μL of 0.5 mM DTNB [Sigma Aldrich, USA (D8130)] in phosphate buffer. The remaining well for each sample was used as blanks and loaded with 200 μL of phosphate buffer only. The plate was then incubated in the dark for 30 min at 22 °C before absorbance was read at 412 nm using a plate reader with correction for the blank reading. Finally, the concentration of plasma thiol level was determined by comparison with a standard curve generated under identical experimental conditions and normalised to the corresponding protein level in the sample and expressed as units/mg homogenate protein.

### 2.10 Thio-Glo Assay

Plasma thiol levels were detected via a Thio-Glo assay based on published protocols [31]. A 24 mM stock solution of ThioGlo^®^ 1 fluorescent thiol reagent [Berry & Associates, USA (HC9080)] was prepared in acetonitrile and stored in dark at 4 °C. Immediately prior to use, the stock was further diluted in PBS (1:100 v/v) to yield the working solution. In a clear, flat-bottomed 96-well plate [Sigma Aldrich, USA (3599)], 5 μL plasma samples were added to wells in duplicates all and made up to 50 μL with nanopure water. Standards were prepared by diluting a 0.5 mM stock solution of 7.7 mg GSH [Sigma Aldrich, USA (G4251)] in nanopure water to give GSH standards ranging from 0 to 25 μM with final volume made up to 50 μL. Next, the diluted ThioGlo solution (50 μL) was then added to all wells, samples were mixed for 10 min and incubated for 5 min in the dark at 22 °C in a plate reader before fluorescence was measured at excitation 384 nm and emission at 513 nm. Plasma thiol concentration was then calculated by comparison to the standard curve generated under identical experimental conditions and normalised to the corresponding protein level in the sample and expressed as μM/ug homogenate protein.

### 2.11 GSH/GSSG assessment

Colonic glutathione (GSH) levels were measured in samples of colon homgenate by optical absorbance at 412 nm following reaction with DTNB: glutathione reductase (GR) solution in 1:1 ratio and β-NADPH [33]. DTNB [Sigma Aldrich, USA (D-8130) (6 mg) and 80 μL of glutathione reductase [GR, Sigma Aldrich, USA (G3664) were prepared fresh in 9 mL and 6 mL of 0.1M Potassium phosphate EDTA buffer, respectively, and the plates incubated (30 s at 22 °C). Next, 2.5 mg β-NADPH [Sigma Aldrich, USA (N-7505)] was diluted in 7.5 mL of KPE immediately prior to reading and 60 μL was added to each well. For the detection of glutathione disulfide (GSSG), samples were pretreated with 10% v/v triethanolamine [TEAM, Sigma Aldrich, USA (T1377)] then incubated with 10% v/v 2-vinylpyridine [Sigma Aldrich, USA (132293)] for 60 min prior to reaction with DTNB:GR solution and β-NADPH. GSH and GSSG standards were prepared from serially diluted stock solutions of powdered GSH [Sigma Aldrich, USA (G-4251)] and GSSG [Sigma Aldrich, USA (G4376)]. Where required, absorbance was immediately measured at 412 every 30 s for 5 min. The change in absorbance per minute (slope) was calculated for each sample and standard in a GSH and GSSG assay. Colonic concentrations of GSH and GSSG were then determined by comparison to the standard curve generated under identical experimental conditions. Finally, GSH was normalised against total glutathione (GSH + GSSG) to give a GSH/GSSG ratio, which is indicative of the glutathione redox status in colonic samples.

### 2.12 Histological analysis

Hematoxylin and eosin (H&E) and Alcian Blue/Safranin O stained tissue sections (10 μm thickness) were imaged over 20-40 fields of view per sample using Zen 2 (Blue edition) imaging software. Histological assessment of colonic structure on H&E stained colon sections was scored by a single, blinded operator based on the following criteria: crypt structure, leukocyte infiltration, epithelium integrity, goblet cell count and edema [34] (histological criteria described in detail in Supplementary Table 1). Each criterion was scored from 0 (no damage) to 2 (severe damage), creating a total score out of 10 for each field of view. A summation of total scores was then normalised against the number of fields of view across the entire length of the colon to provide an average histological score. Positive mucin staining using Alcian blue and Safranin O stained colon sections were quantified using Metamorph^®^ imaging analysis software (*V 7.6*) and standardised over the length and area of the imaged colon, respectively.

### 2.13 Alcian Blue and Safranin O staining

Colon sections were deparaffinised, hydrated and placed into glass Coplin staining jars filled with Alcian Blue solution (adjusted to pH 2.5) and incubated at 22 °C for 30 min. The slides were then rinsed in distilled water and counterstained in 0.1 % (w/v) acetic safranin solution for 5 min. Following this, the slides were washed, subsequently dehydrated through a graded alcohol series before cleared in xylene and cover-slipped with DPX [Sigma-Aldrich, USA (06522)].

### 2.14 Colon length

Excised colons were imaged against a precise ruler, and the length of the colon was determined by obtaining the pixel length of the colon using Metamorph^®^ imaging analysis software (V7.6) and normalising to the length of the ruler captured together within each colon image. The value obtained reflected the number of Pixels/ cm of physical length of colon. These values were then averaged across the cohort of samples, and the average number of Pixels/ cm value was used to reverse calculate the colon length of all individual samples.

### 2.15 TUNEL-based detection of non-viable cells

The level of cell viability in colon sections was measured using a DeadEnd™ Fluorometric TUNEL kit [Promega, USA]. Tissue sections (5 μm) were deparaffinised in xylene, hydrated in graded alcohol and immersed in 0.85% NaCl, and subsequently PBS for 5 min. Slides were then fixed for 15 min in 4% (v/v) formaldehyde. 100 μL of 20 μg/mL Proteinase K solution was then added to each slide for 9.5 min. Slides were then washed in PBS before fixing in 4% (v/v) formaldehyde for 5 min. Equilibration buffer (100 μL) was then added for 10 min and 50 μL of rTDT reaction mix was subsequently added to the tissue area and incubated for 60 min at 37 °C. The reaction was stopped by immersing slides in 2 x SSC buffer for 15 min and rinsing 3 times in PBS, 5 min each. Slides were then counterstained with 50 μL of Spectral DAPI [PerkinElmer, USA (FP1490)] at 1:800 v/v concentrations in 1xTBS-T and subsequently washed in PBS before coverslipped using a DAKO fluorescence mounting agent [Agilent Technology, USA (S3023)].

### 2.16 Immunofluorescence detection of Nrf2 and GClC

Colon sections (n=12 per group; 5 μm thickness) were deparaffinsed in xylene and rehydrated through graded alcohols. Endogenous tissue peroxidase was blocked by incubating slides with 3% (v/v) hydrogen peroxide [Chem-supply, Gilman, South Australia] for 5 min. The slides were then rinsed with 1x TBS-T (3X), blocked with a commercial caesin solution [Dako, USA] and all slides were then incubated for a further 30 min at 22 °C. Next, slides were incubated with 50 μL of unconjugated polyclonal primary antibody (anti-GCLC 1: 200 v/v, anti-Nrf2 1: 200 v/v, Abcam) for 1 h at 22 °C and rinsed with 1x TBS-T. Treated sections were then incubated with secondary 1:200 v/v anti-rabbit horseradish peroxidase for 30 min at 22 °C. After rinsing three times with 1x TBS-T, slides were finally incubated in 1:50 v/v tyramide signal amplification Opal520 fluorophore [PerkinElmer, Massachusetts, USA] for 10 min and washed in 1x TBS-T. Specimens were then counterstained with 1:1000 v/v Spectral DAPI [Perkin Elmer, Massachusetts, USA] for 5 min and washed three times with 1x TBS-T before being mounted with fluorescent mounting medium [Dako, USA]. Where required slides were scanned using Zeiss AxioScan.Z1 at 200x magnification. Metamorph^®^ imaging analysis software (V 7.6) was then used to quantify the extent of TUNEL/Nrf2/GClC immuno-positive fluorescent staining and standardised over the entire length of the colon specimen.

### 2.17 Statistical analysis

All statistical analysis was performed with GraphPad Prism^®^ (V 7.02) using one-way analysis of variance (ANOVA) with Tukey’s correction, except with mice behavioural data whereby two-way analysis of variance (ANOVA) with Tukey’s correction was employed. Data represented as mean ± standard deviation and statistical significance was established at *p* <0.05.

## 3. Results

### 3.1 Acclimation of mice to water additives

To ensure that mice from each treatment group received similar levels of salt in drinking water, behaviours associated with water-seeking and drinking in the Intellicage were quantified in individual animals. Behaviour was also examined to ascertain what, if any, impact of each treatment was evident on water-seeking activity. From 0 to 2 days during DSS challenge, mice from all four treatment groups made similar numbers of visits to their allocated corners suggesting similar levels of water consumption and tolerance to NaCl/NaSCN salts added into normal and DSS-supplemented drinking water. Hence, this was a period where insult with DSS did not alter behaviour of mice allocated to develop experimental colitis. Mice from all four treatment groups made similar numbers of visits to their allocated corners indicating general exploratory activity was similar and were tolerant to NaCl/NaSCN salts added into normal and DSS-supplemented drinking water.

Progression of disease for mice in DSS-supplemented groups resulted in diminished total number of visits to their allocated corners. For example, mice treated with DSS alone showed a 6-fold reduction in all water-seeking behavioural parameters (Figure 1A-D) (compare values for Day 0 vs 7; p < 0.001), while the corresponding salt-supplemented DSS-control group showed a 4-fold reduction in seeking water (compare salt control vs DSS+Cl^-^ groups at Day 7; *p* < 0.001) consistent with the development of acute colitis. Similarly, mice co-supplemented with NaSCN together with DSS in the drinking water (DSS/SCN^-^) showed a 5-fold reduction in all water-seeking behavioural parameters (compare salt control vs DSS+SCN^-^; *p* < 0.001). In the absence of DSS, mice showed similar drinking behaviour with total visits largely unchanged compared to the corresponding day 0 values irrespective of the presence or absence of added salt (Figure 1 A). Although this is most likely due to disease progression, conditioned taste aversion could also be a contributing factor, whereby the association sickness with a particular location, food, fluid etc results in a learned aversion.

**Figure 1.**
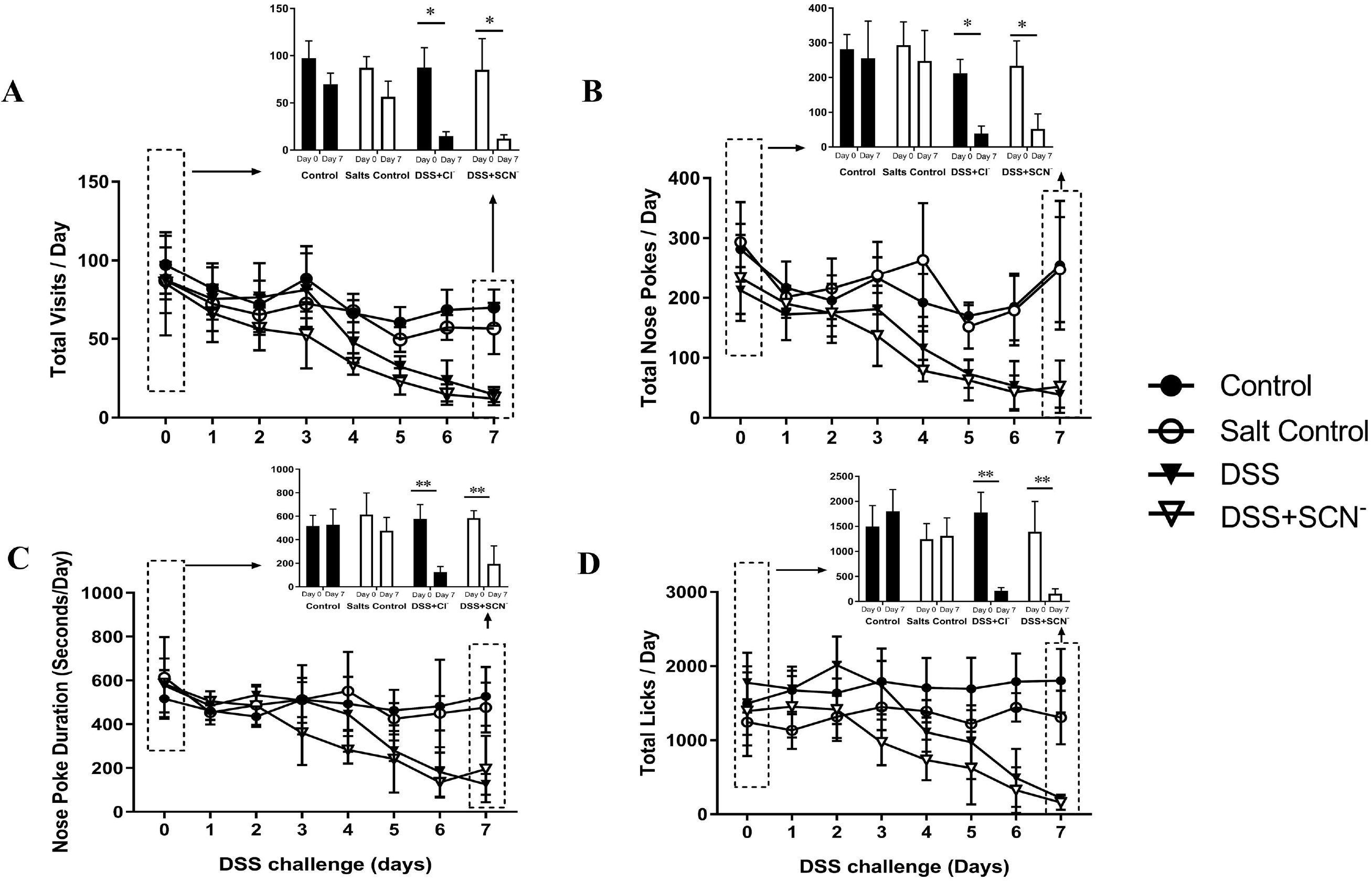
Acclimation to water additives and measurement of behavioural parameters for mice supplemented with or without NaSCN in drinking water during 8 days of DSS challenge. Female C57BL/6 mice (9 weeks of age) were allocated to groups, individually tagged with an RFID transponder and trained to visit and nose poke to obtain water located at designated corners of the IntelliCage system over 2-3 weeks. Once adequately trained to seek water at the designated corner, all mice were continuously monitored for 8 days during the active phase of experimental colitis. The number and duration of visits, licks and nose pokes for each mouse were automatically recorded and behavior quantified. Panels show (A) Total visits to allocated drinking corners per day, (B) Total nose-pokes to allocated drinking corners (C) Nose poke duration and (D) Total licks. Data represents mean ± SD; n=6 per group. * Different to the corresponding day 0 value in the same group; \**p* < 0.001 or ** *p* < 0.0001. # Different to the control group; *p* < 0.001. ## Different to the salts control group; *p* < 0.001.

The monitoring of water seeking (described above) was representative of other behavioural parameters monitored including visits, nose pokes, nose poke duration and total licks, likely reflecting a reduction in fluid intake colitis severity progressed. No statistical difference in any clinical parameters was observed between the NaSCN supplemented DSS group and NaCl supplemented DSS group (Figure 1 AD).

### 3.2 Monitoring clinical markers of experimental colitis

Mice provided DSS in the drinking water showed a significantly greater weight loss when compared to both the mean at day 0 (*p* < 0.001) and the average weights determined from the corresponding control groups at day 7 (*p* < 0.001). Notably, mice co-supplemented with DSS/SCN^-^ in the drinking water showed a significantly higher weight loss than mice supplemented with DSS + Cl^-^ (*p* < 0.05) (Figure 2A), suggesting that SCN^-^ did not generally inhibit the loss of body condition in these mice, but possibly worsened the colitis. Overall, DSS-challenged mice exhibited significantly shorter colons in comparison to mice from corresponding water and salt controls (~1.4-fold shorter colon; *p* < 0.001). However, no difference in colon length was observed between mice co-supplemented with either DSS/SCN^-^ or DSS/Cl^-^ in the drinking water (Figure 2B).

**Figure 2.**
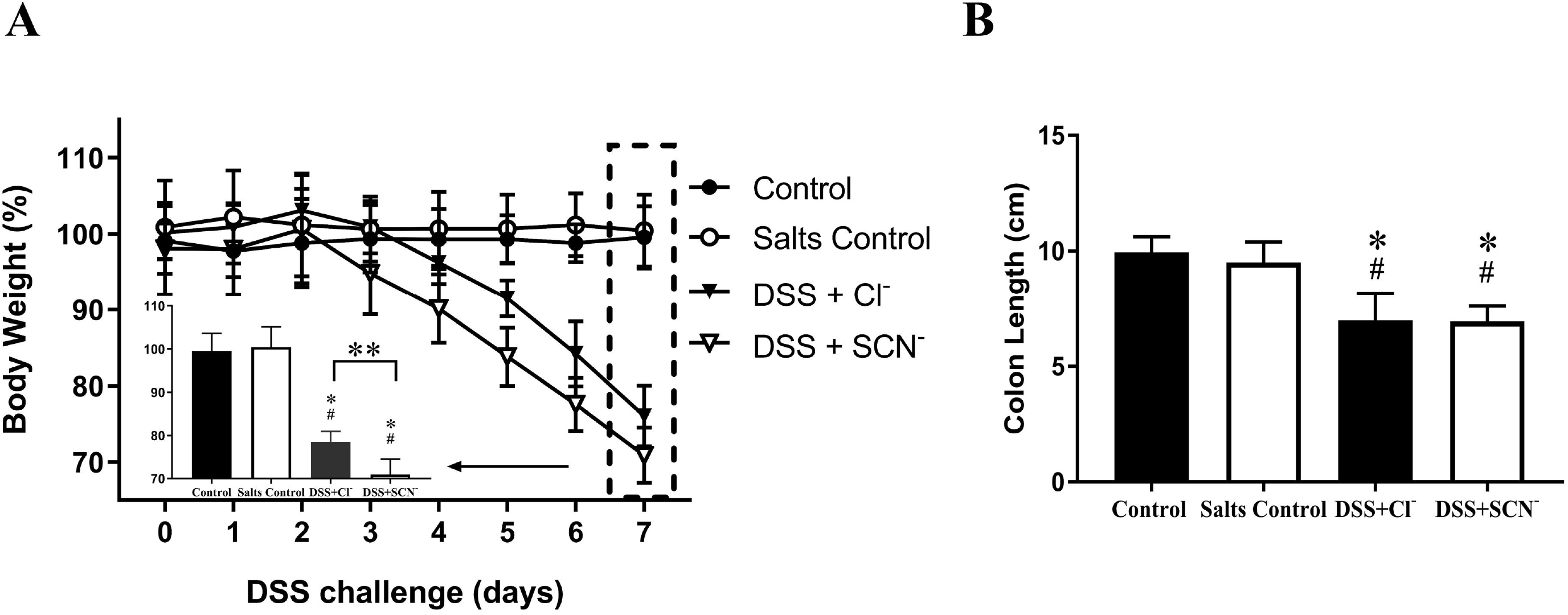
Clinical and physiological markers for mice over 8 days of DSS Challenge. Female C57BL/6 mice (9 weeks old) were treated with water (control), water + 0.7 mM NaCl + 10 mM NaSCN (salts control), 3% w/v DSS + 10 mM NaCl (DSS+Cl^-^) and 3% w/v DSS + 10mM NaSCN (DSS +SCN^-^) over 8 days. Panel (A) Weight loss calculated as a percentage of the mean body weight at day 0 of the control groups before treatment commenced; (B) Colon length calculated using Metamorph^®^ computer analysis system (V 7.1). Data represents mean ± SD; n=6 per group. * Different to control; *p* <0.001. ^#^Different to salts controls; *p* < 0.001. ** Different to DSS-Cl^-^ mice; *p* < 0.05. Note, in some comparisons between DSS/Cl^-^ vs DSS/SCN^-^ groups a near statistical outcome was achieved; *p* = 0.051.

### 3.3 Tissue and plasma biochemistry

#### 3.3.1 Tissue and circulating levels of SCN^-^

As anticipated, mice receiving SCN^-^ in water exhibited higher levels of fecal SCN^-^ than the corresponding salt control (*p* < 0.01). However, while the fecal level of SCN^-^ increased in animals co-supplemented DSS/SCN^-^, this was not statistically different to the corresponding DSS/Cl^-^ group. Nevertheless, mice co-supplemented DSS/SCN^-^ achieved a ~3-fold increase in fecal SCN^-^ in comparison to their DSS/Cl^-^ counterparts (Figure 3A).

**Figure 3.**
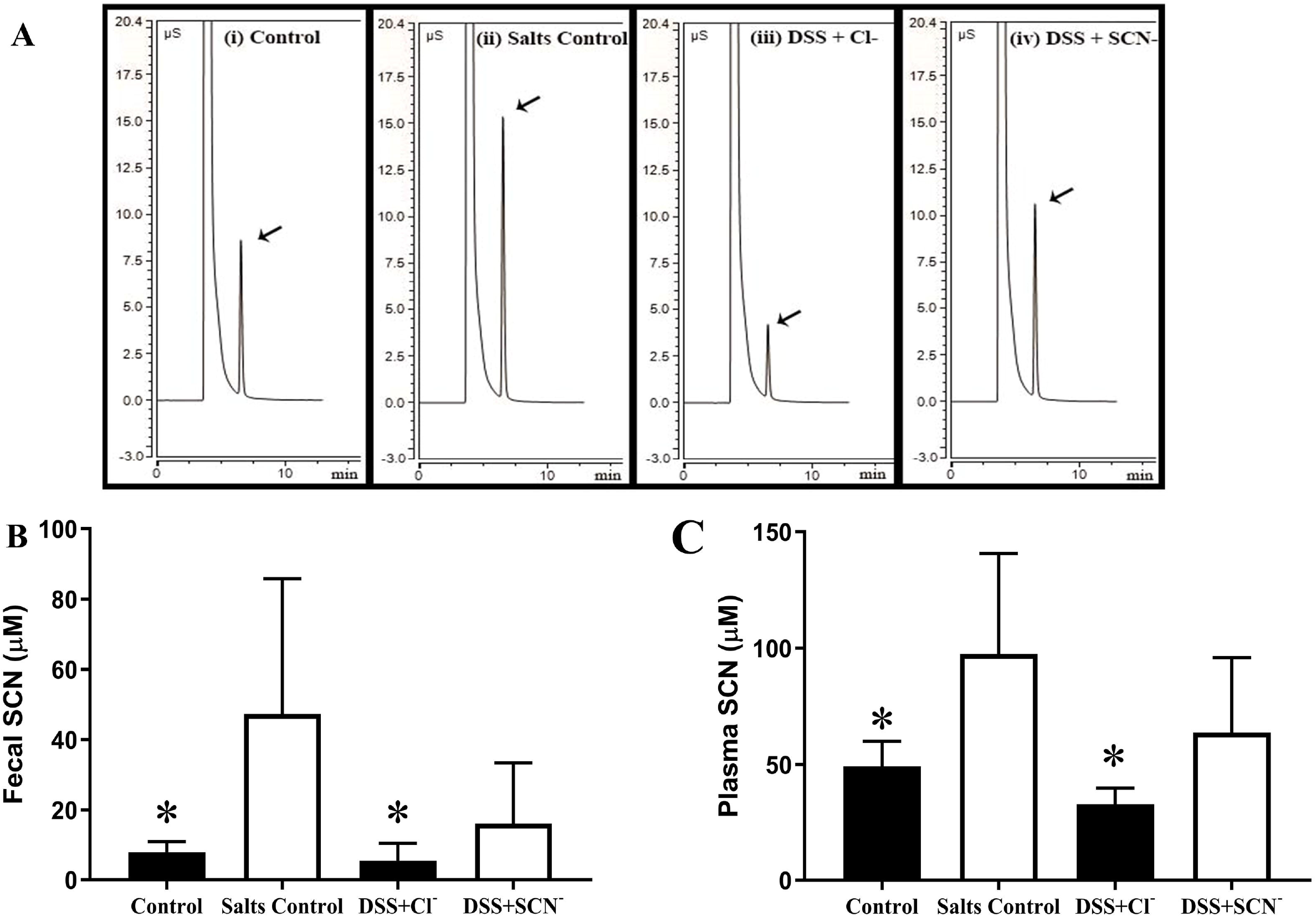
Plasma and fecal levels of SCN^-^ measured in DSS-treated mice with or without SCN^-^ supplemented in drinking water. Fecal samples were obtained 2 days after initiating the study, the isolated pellets were homogenised and stored at −80 °C. Next, plasma was isolated from mice at completion of the study (day 7). Separate samples of fecal homogenate and plasma were mixed 1:1 v/v with acetonitrile, centrifuged and SCN^-^ levels were measured by ion-exchange chromatography and quantified using Chromeleon data system software (*v.7.1*) using a standard curve prepared with authentic SCN^-^. Panels show representative chromatograms for (A) plasma SCN^-^ (arrow) and quantified levels of SCN^-^ in (B) fecal and (C) plasma samples. Data represents mean ± SD; n=6 per group. * Different to salts control; *p* < 0.05.

In the absence of DSS, plasma SCN^-^ levels in the salt control mice increased markedly to 0.1 mM in comparison to the corresponding control (*p* < 0.05). However, in the presence of DSS, the plasma SCN^-^ concentration was lower, reaching ~0.07 mM, whereas mice from the DSS/Cl^-^ group had approximately 0.04 mM serum SCN^-^ levels. The decrease in serum SCN^-^ in mice supplemented with NaSCN could be explained by the significant reduction in fluid intake during the course of DSS-induced colitis (Figure 1). In addition, malabsorption from acute severe diarrhoea experienced during DSS – induced colitis is likely to contribute to a decrease in serum SCN^-^ from dietary or supplemented sources (Figure 3B). Assessment of colon tissue levels of SCN^-^ was unable to be performed due to interference from complex co-eluting, overlapping peak responses.

#### 3.3.2 Ex vivo competitive inhibition of MPO-mediated luminol oxidation

We have previously demonstrated that luminol is preferentially oxidised by the two-electron oxidant, HOCl – produced chiefly by MPO in the presence of free Cl^-^ and H_2_O_2_ [30]. Peroxidase-mediated oxidation of luminol substantially increases the luminescent intensity of luminol [35].

Herein, we attempted to measure the *ex vivo* inhibitory effect of SCN on the hypocholorous activity of purified human MPO in the presence of free Cl^-^ and H_2_O_2_. We observed a significant dose-dependent decrease in HOCl-mediated oxidation of luminol with increasing concentrations of NaSCN (Figure 4). A 60% decrease in MPO-mediated luminol oxidation was observed at 0.1 mM NaSCN, while luminol oxidation was almost abrogated (>95%) at NaSCN concentrations over 1 mM. The IC_50_ value (best-fit) was 0.034 mM NaSCN (Figure 4; insert).

**Figure 4.**
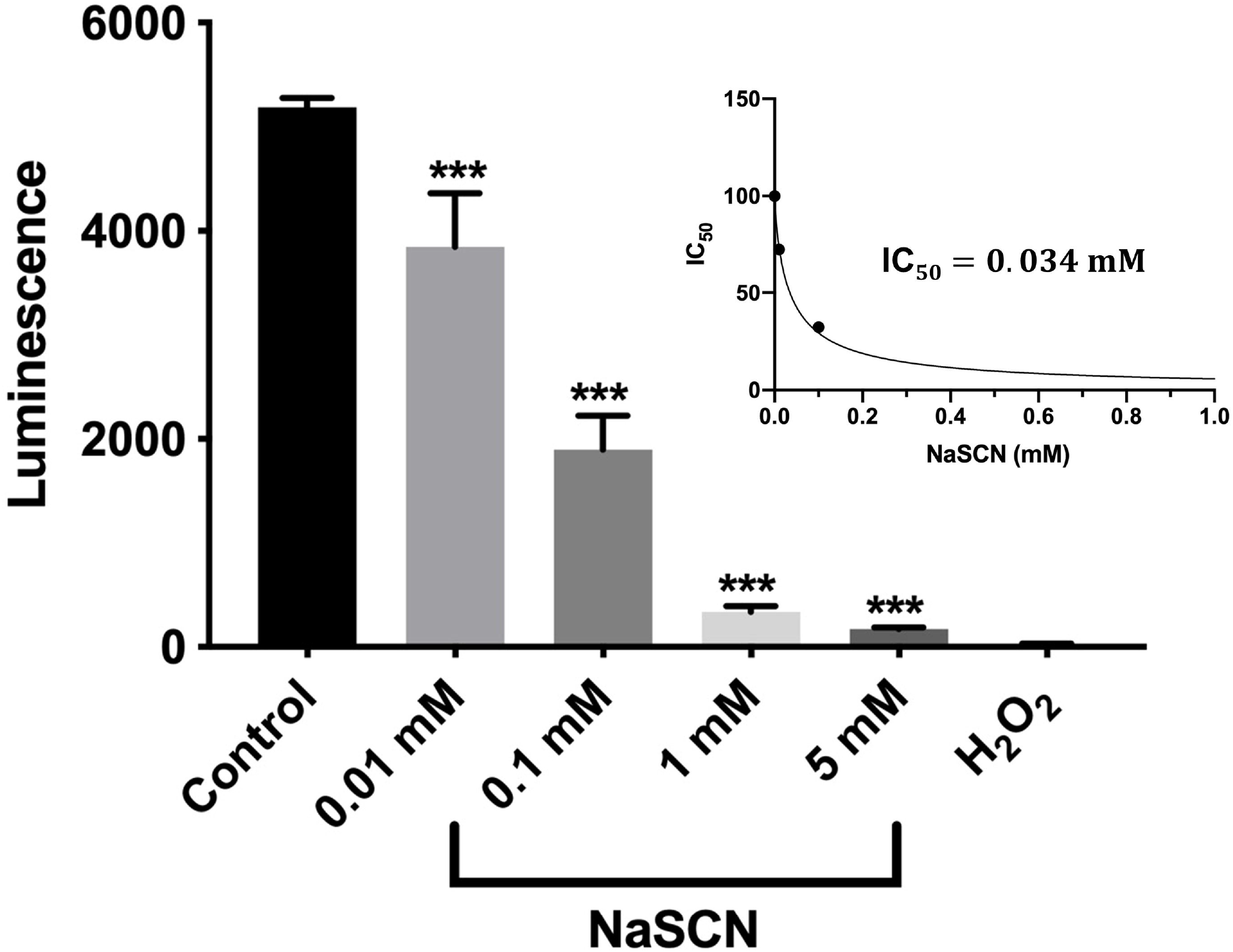
*Ex vivo* SCN^-^-induced competitive inhibition of myeloperoxidase. Ranging concentrations of sodium thiocyanate (NaSCN; 0.01 mM, 0.1 mM, 1 mM and 5 mM) in a mixture to a final concentration of 0.2 μg/mL myeloperoxidase (MPO), 67 mM NaCl, 0.8 mM luminol and 45 mM H_2_O_2_ inhibited oxidation of luminol in a dose-dependent manner. Control group was devoid of NaSCN, while H_2_O_2_ alone group was devoid of NaSCN and MPO. Insert: Best-fit IC_50_ value determined from concentrations of 0 mM, 0.01 mM, 0.1 mM and 1 mM of NaSCN. Experiment was repeated n=5 in duplicates. Data represented as mean +/- SD. ***Different to control group; *p* < 0.001.

#### 3.3.3 Assessing neutrophil activity in the colon

Quantitative ELISA revealed that mice supplemented with DSS/Cl^-^ or DSS/SCN^-^ showed significantly elevated levels of calprotectin in the colon tissue relative to the corresponding controls, consistent with DSS stimulating cellular infiltration to the gut tissues (Figure 5A). However, no difference in calprotectin was determined between groups of mice co-administered DSS/SCN^-^ or DSS/Cl^-^, indicating that SCN^-^-treatment had no impact on the degree and/or the extent of neutrophil recruitment and activation (Figure 5A).

**Figure 5.**
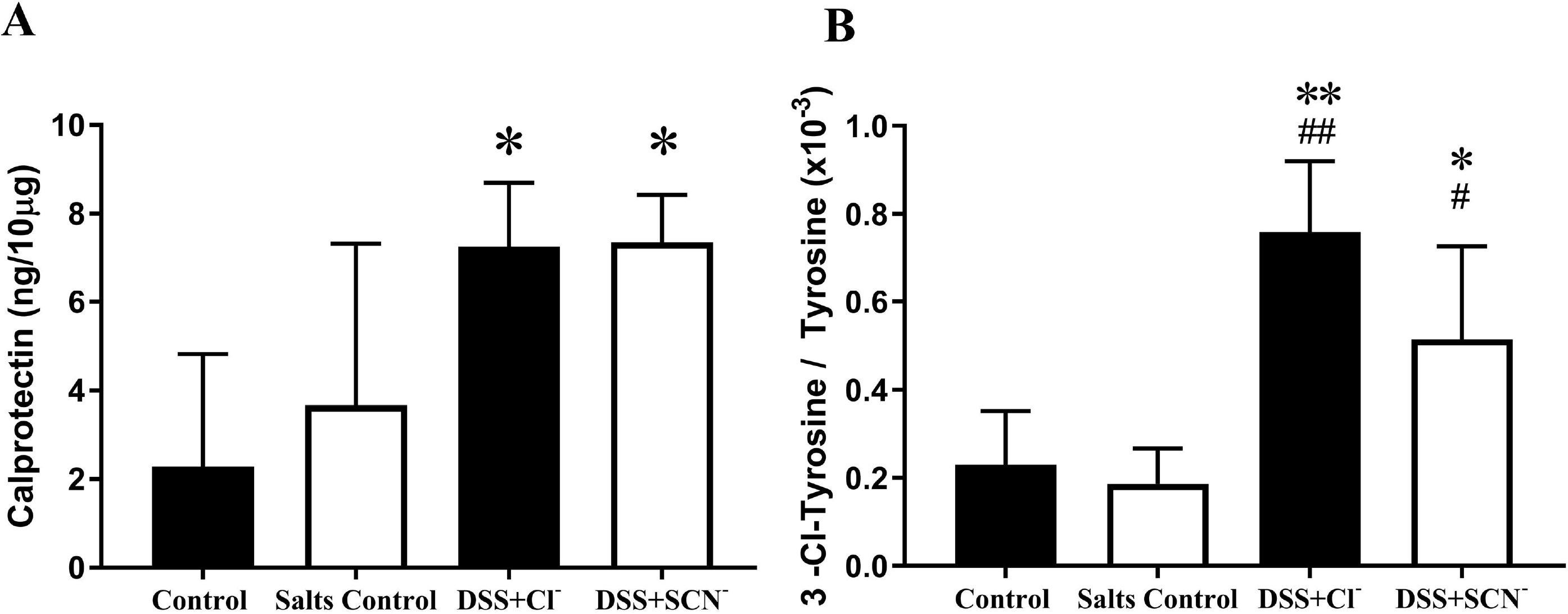
ELISA detection of calprotectin and LC-MS measurement of 3-Cl-Tyr, a specific marker for HOCl-mediated oxidative damage in colon homogenates. Panel (A) shows levels of calprotectin in colon homogenates measured via ELISA. Levels of 3-Cl-Tyrosine and Tyrosine were assessed in colon homogenates with liquid chromatography coupled with quantitative mass spectrometry using labelled 3-Cl-Tyrosine and Tyrosine as internal standards, respectively. Ion quantification was performed and 3-Cl-Tyrosine to tyrosine ratio was determined, shown in panel (B). Note in (B) DSS + Cl^-^ vs DSS + SCN^-^ *p* = 0.051. Data are shown as mean ± SD; n=6 per group. * Different to control; \**p* < 0.05 or \*\**p* < 0.001. ^#^Different to the salts control; ^#^ *p* < 0.05, ^##^ *p* < 0.001.

Following the investigation of colonic calprotectin as a surrogate biomarker for neutrophil infiltration/activation, the potential for SCN^-^ to redirect the pattern of neutrophil-MPO-mediated oxidative damage was examined. As demonstrated (Figure 3B & C), fecal and plasma SCN^-^ levels increased in mice supplemented with thiocyanate; therefore, it is feasible that MPO production of HOCl may be shifted to HOSCN if SCN^-^ can compete with the pool of Cl^-^ for enzymic halogenation in the colon. Any shift toward production of HOSCN may impact on the level of oxidation of biological targets as HOSCN shows greater selectivity toward reduced thiols. The oxidation product 3-Cl-Tyrosine is a specific biomarker for HOCl-mediated protein oxidative damage [31]. Hence the assessment of 3-Cl-Tyrosine can reveal the degree of HOCl involvement in oxidative tissue damage and its level of production in animals supplemented DSS/Cl^-^ and DSS/SCN^-^.

Quantification of the LC-MS outcomes indicated that the DSS-supplemented mice showed a higher 3-Cl-Tyrosine/Tyrosine ratio than corresponding controls, consistent with the recruitment and activation of neutrophils to the inflamed colon (as confirmed here by measuring colonic calprotectin levels, Figure 5A). Notably, the colonic 3-Cl-Tyrosine/Tyrosine ratio was decreased in mice co-supplemented DSS/SCN^-^ compared to mice treated with DSS/Cl^-^ although, this diminution was not statistically significant (*p*<0.051). Overall, thiocyanate appeared to divert HOCl-mediated oxidative damage away from protein tyrosine residues in DSS challenged mice (Figure 5B). Note, representative LC-MS spectra shown in Supplementary data (Figure S1).

#### 3.3.4 Assessing thiol oxidation in the colon

Data sets above demonstrated that the level of neutrophil recruitment and activation in the colon were similar in mice co-supplemented with both DSS/SCN^-^ or DSS/Cl^-^. However, co-supplementation of SCN^-^ in DSS-challenged mice may have diverted neutrophil-MPO to produce HOSCN, which has been previously demonstrated to preferentially oxidise thiol residues [36]. To investigate whether thiol oxidation was selectively enhanced in mice co-supplemented with SCN^-^/DSS, various approaches were used to assess plasma thiol status (Figure 6). As anticipated, mice in the DSS-treatment groups presented with significantly decreased plasma thiol levels in comparison to the corresponding controls (*p* < 0.05) irrespective of whether determined as the plasma thiol redox status (Figure 6A) or the total concentration of plasma thiol determined using two different approaches (Figure 6B and C). Interestingly, serum thiol oxidation was consistently higher when measured by the ThioGo assay, compared to the DTNB approach (Figure 6C & B) and this is likely due to ThioGlo binding non-specifically to plasma proteins in a non-thiol dependent manner. However, no difference in plasma thiol oxidation was determined in samples taken from mice co-supplemented SCN^-^/DSS or Cl^-^/DSS, indicating that SCN^-^ was unlikely to compete effectively with Cl^-^ as the primary substrate for the MPO halogenation cycle, at least when SCN^-^ is present at plasma concentrations that match SCN^-^ levels determined for human smokers.

**Figure 6.**
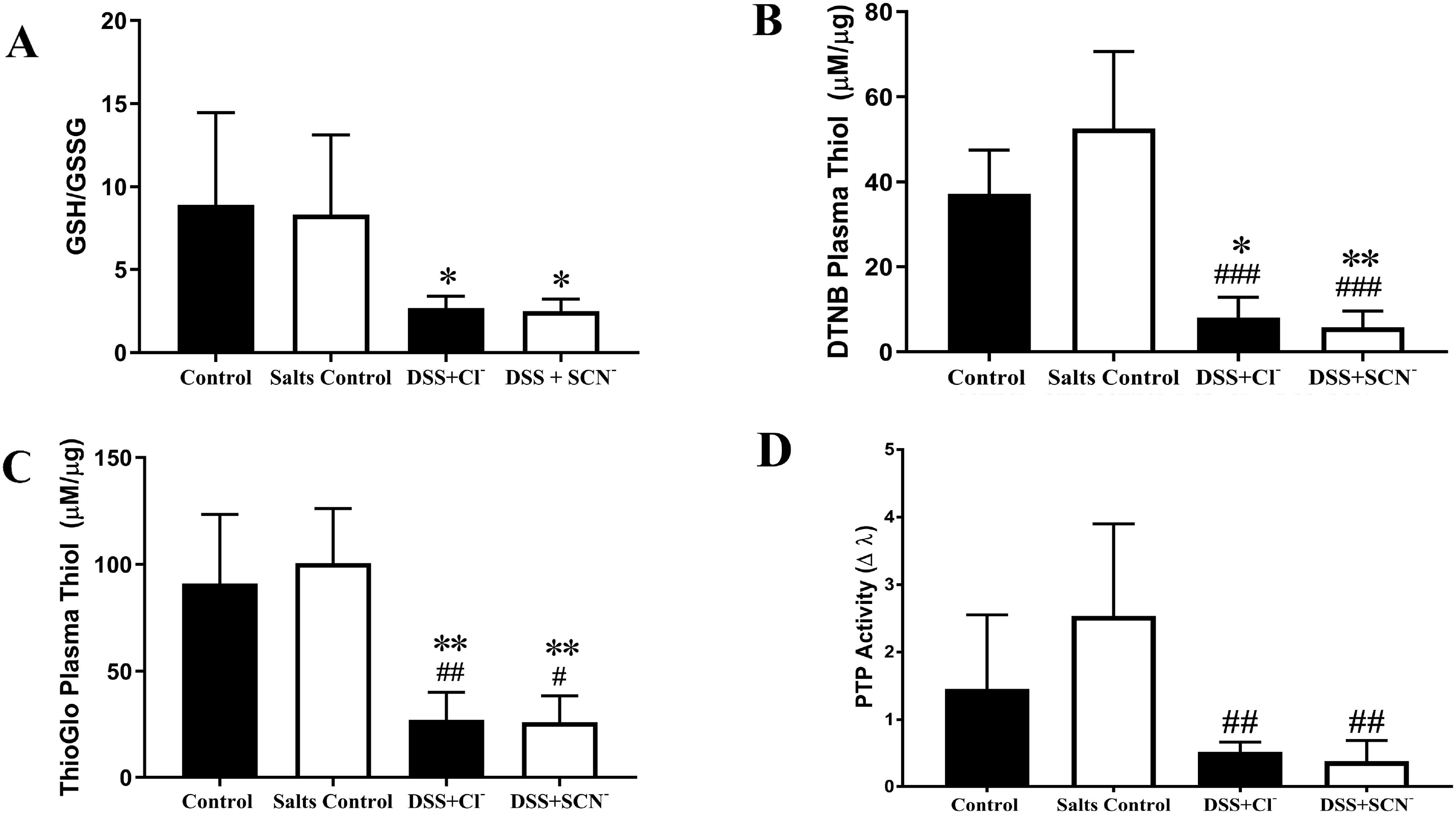
Plasma protein thiol levels and protein tyrosine phosphatase (PTP) activity in mice following 8 days of DSS challenge. Plasma was assessed for content of total thiols and specific low molecular weight thiol (glutathione) levels. PTP levels in plasma samples were measured via reaction with an alkaline and acid phosphatases substrate (pNPP) over 20 hours and absorbance was measured at 405 nm. Panels show (A) plasma glutathione oxidation levels expressed as the ratio of reduced glutathione (GSH) / oxidised glutathione (GSSG); determination of plasma thiol levels with (B) Ellman’s reagent (DTNB) and (C) ThioGlo^®^ 1 (MMBC); (D) PTP activity based on Δ absorbance. Data are mean ± SD; n=6 per group. * Different to Control; \**p* < 0.05, \*\**p* < 0.005, \*\*\**p* < 0.001. ^#^ Different to Salts Control; ^#^*p* < 0.05; ^##^*p* < 0.005, ^###^ *p* < 0.001.

Next, we tested whether SCN^-^ altered colon tissue thiol oxidation levels. The family of protein tyrosine phosphatase (PTP) enzymes contain a conserved protein thiol group at the active site; this cysteine residue is susceptible to oxidation yielding inactive PTP [37–40]. Thus, PTP activity is a considered to be a sensitive surrogate marker for tissue thiol oxidation status. Assessment of total plasma PTP revealed significantly lower PTP activity in both DSS-treatment groups relative to the corresponding control (*p* < 0.05) (Figure 6D), reflecting the measurements of plasma thiol content. However, similar to the case in plasma, PTP activity in colon homogenates did not differ between mice co-supplemented DSS/SCN^-^ or DSS/Cl^-^ (Figure 6D). Taken together, this data indicates that the addition of SCN^-^ to DSS-challenged mice does not influence total levels of thiol damage in plasma or colon tissues.

### 3.5 Immunohistochemical assessment of colon protein expression

Our data indicated that thiol oxidation in DSS-stimulated mice remained similar irrespective of the presence or absence of SCN^-^. We considered that MPO-generated HOSCN may not preferentially target thiols or that thiol turnover may be altered following SCN^-^ supplementation. The transcription factor nuclear factor erythroid 2-related factor 2 (Nrf2) is a master regulator for many antioxidant genes and also encodes for glutamate-cysteine ligase catalytic subunit (GCLC), the rate-limiting enzyme in the production of the low molecular weight thiol GSH [41–43]. Therefore, to investigate whether a change in thiol production may have some bearing on thiol levels determined in the plasma and colons of mice in the DSS/SCN^-^ or DSS/Cl^-^-treatment groups, the Nrf2-GCLC axis was investigated in the colons of mice. Immunohistochemical staining of Nrf2 revealed an intense, transmural pattern of colonic Nrf2 expression across both DSS treatment groups, while only moderate Nrf2^+^-staining was observed in the corresponding control groups; where present Nrf2^+^-immune fluorescence staining was primarily localised to colon epithelial cells in these control tissues (Figure 7A).

**Figure 7.**
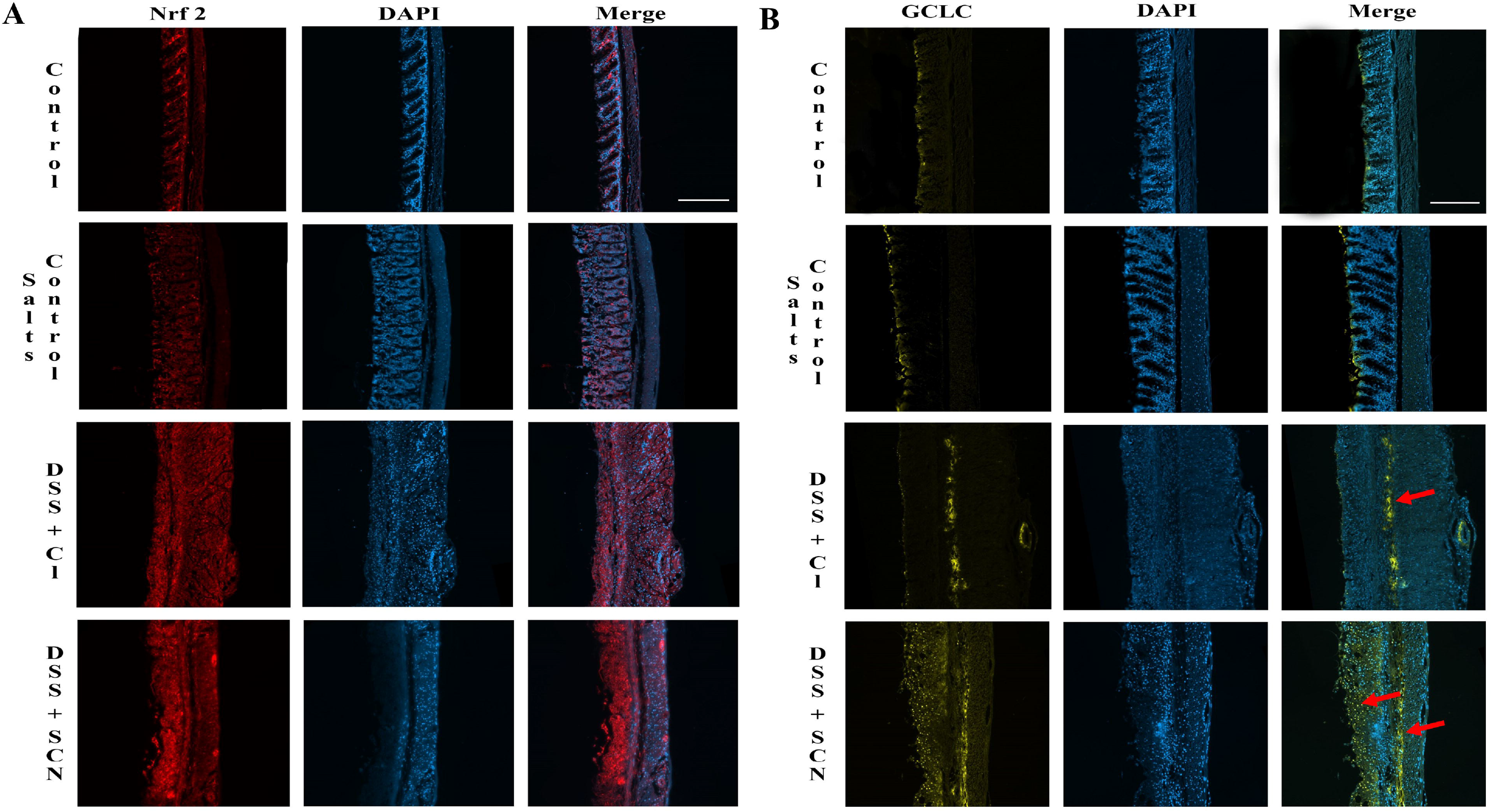
Immunofluorescence-based analysis of antioxidant production via Nrf2 and GCLC labelling. **Mice** colon sections were labelled with Nrf2 and GCLC. Images were captured using the Zeiss AxioScan at 10x magnification. Panels show representative images of (A) Nrf2 (red) and nuclei (blue, DAPI) (B) GCLC (yellow, red arrow) and nuclei (blue, DAPI) in inflamed colon together with merged/co-registered images. Scale bar = 100 μm.

Parallel GCLC^+^-immune fluorescence staining revealed scarce GCLC expression in both control mouse groups, which contrasted markedly in comparison to the higher levels of GCLC^+^-immune fluorescence staining in colons from mice challenged with DSS. Patterns of GCLC expression were also different between the control and DSS-treated cohorts. Similar to the corresponding expression pattern for Nrf2^+^- immune fluorescence, GCLC^+^-immune fluorescence was generally restricted to the epithelial lining in the colon specimens obtained from mice in the control groups. By contrast, mice exposed to DSS in the drinking water showed more GCLC^+^-immune fluorescence in the submucosa and in particular, mice co-supplemented with DSS/SCN^-^ showed a marked increase in GCLC protein expression in the colon submucosa with GCLC^+^-immune fluorescence staining distributed in a clear punctate pattern throughout the tissue (fB).

Subsequent quantification of Nrf2^+^-immune fluorescence labelling confirmed significantly higher levels of Nrf2 in DSS treatment groups relative to the controls (*p* < 0.05). Also, Nrf2^+^ expression was near identical between the DSS/SCN^-^- and DSS/Cl^-^-treatment groups (Figure 8). Numeration of GCLC^+^ pixels revealed a significant increase of GCLC in DSS challenged mice relative to the controls (*p* < 0.001). Moreover, GCLC expression was markedly higher in mice co-supplemented DSS/SCN^-^ relative to mice co-supplemented DSS/Cl^-^ (*p* < 0.05) (Figure 8). Overall, these immune-histochemical data suggest that a selective upregulation of thiol synthesis occurs in mice co-supplemented with DSS/SCN^-^ and that this outcome may add complexity to the interpretation of thiol oxidation studies presented earlier (Figure 8), since the flux of thiol synthesis is markedly different when comparing control vs DSS-treated mice and appears to be altered further in mice assigned to the DSS/SCN^-^ and DSS/Cl^-^ groups.

**Figure 8.**
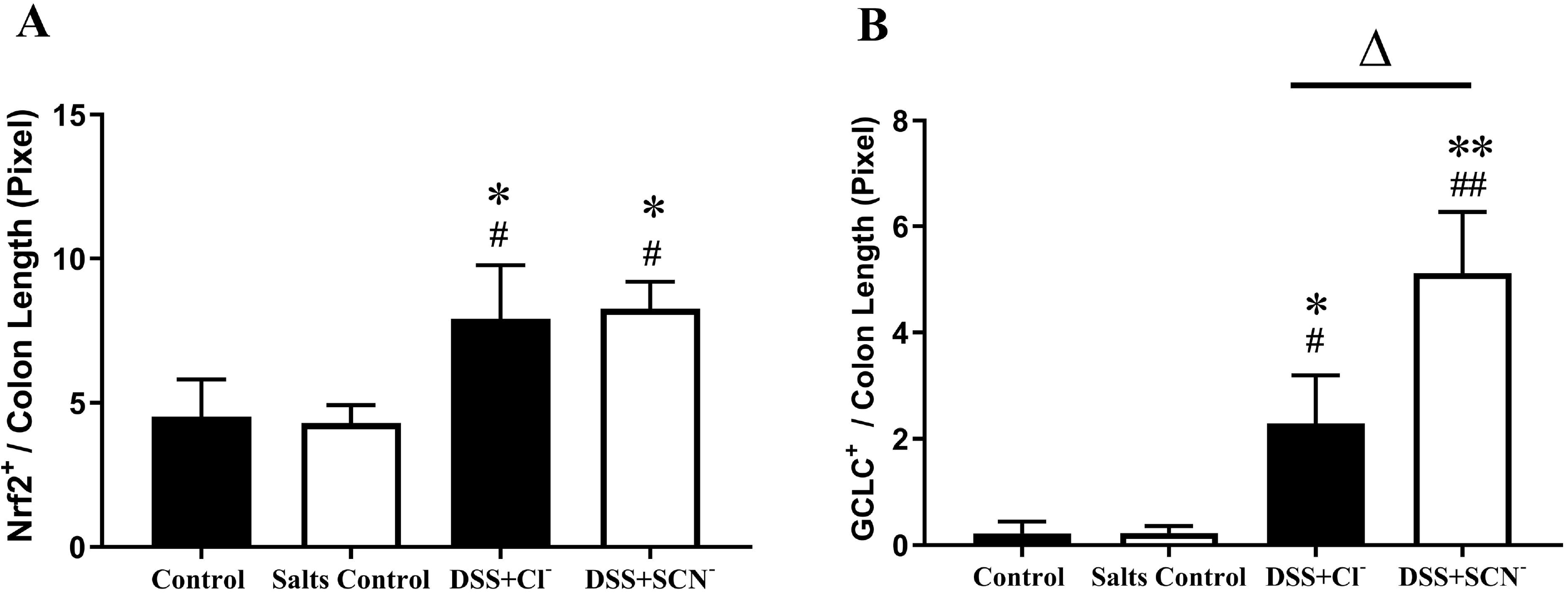
Quantification of Nrf2 and GCLC protein in descending colon sections. Mice colon sections were labelled with (A) Nrf2 and (B) GCLC. Images were captured using the Zeiss AxioScan at 10x magnification. Analysis of Nrf2 immunofluorescence staining in colon sections were quantified and adjusted to corresponding pixel length using Metamorph^®^ V 7.8 software. Data are shown as mean ± SD, n=3 per group. ^Δ^*p* < 0.05 compared to DSS+Cl^-^; *Compared to control; \**p* < 0.05, \*\**p* <0.001; ^#^ Compared to salts control; ^#^*p*<0.05, ^##^*p*<0.001.

### 3.6 Histopathological evaluation of colon damage

Sections of mouse colon from control and salts control groups showed minimal or no signs of histopathology in the transverse and descending colon with relatively intact columnar epithelium and evenly spaced crypts (*e.g*., refer to Figure 9A, upper panels). Evidence of major cellular infiltration, goblet cell loss and oedema were clearly absent in these control tissues. In contrast, DSS-challenged mice presented with severe inflammation and histopathological damage (Figure 9A, lower panels). Cellular infiltration was evident at the mucosal, lamina propria and submucosal layers and at some colon regions a transmural (all layers) pattern of leukocyte inflammation was detected. Mucosal injury including loss of crypts of Lieberkühn was also evident, together with partial to severe distortion of the mucosal epithelial lining. Submucosal and lamina propria-to-transmural oedema was also detected and caused a marked expansion of the intestinal submucosa relative to the control groups. However, no clear differences in the level of histopathological damage were noted between colons isolated from mice assigned to the DSS/Cl^-^ and DSS/SCN^-^ groups (Figure 9A). Subsequent histopathological scoring of the transverse and descending colon revealed similarly high histopathology scores of colons isolated from mice assigned to the DSS/Cl^-^ and DSS/SCN^-^ groups, indicative of severe colonic pathology (Figure 9B).

**Figure 9.**
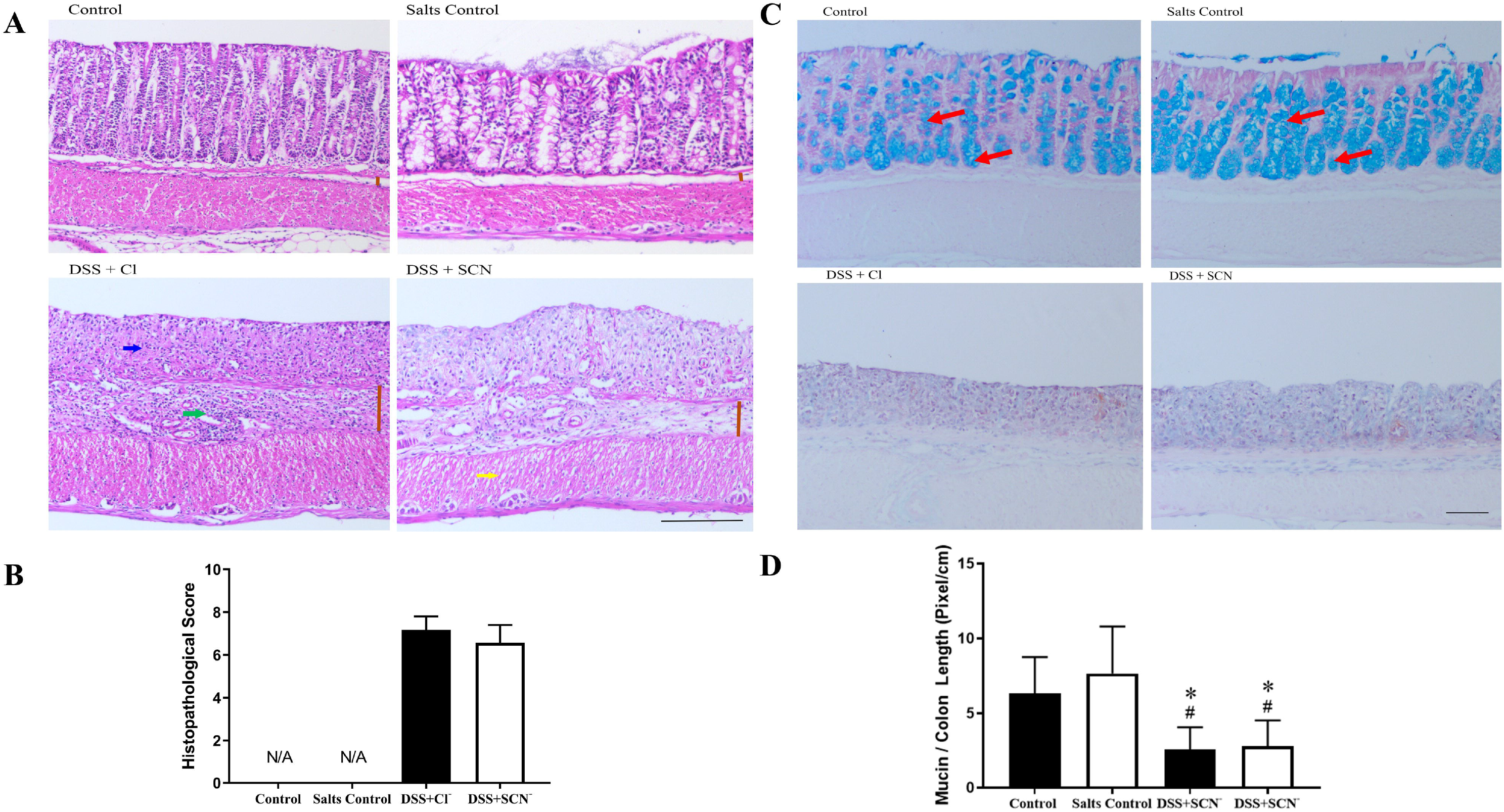
Histopathological examination of colon sections in mice with DSS-induced colitis. (A) Representative haematoxylin and eosin stained colon sections were taken at 10x magnification. Arrows indicate cellular infiltration (green), ulceration (blue) and oedema (yellow). Expansion of intestinal submucosa due to oedema and cellular infiltration are highlighted by comparing the vertical lines (brown). (B) Histopathological scores of transverse and descending colon, based on crypt structure, epithelium integrity, edema, goblet cell levels and leukocyte infiltration. 0 (no pathology), 5 (moderate pathology), 10 (severe pathology). (C) Representative Alcian blue (mucin) and Safranin O (nuclei) staining of colon sections taken at 10x magnification. Arrows (red) indicate areas of mucin aggregation. (D) Quantification of mucin levels in transverse and descending colon determined using Metamorph^®^ (V 7.8) software with staining output normalised to corresponding colon length. All data represents mean ± SD over 25 fields of view per mouse, n=6 per group. * Different to Control; * *p* < 0.005 ^#^ Different to Salts Control; ^#^ *p* < 0.05.

Alcian blue staining of mucin-containing goblet cells in colonic sections revealed that unchallenged (controls) mice displayed minimal-to-no pathology with normal mucin production, indicative of normal goblet cell function (Figure 9C, compare upper panels). In contrast, DSS-treated mice displayed severe to complete destruction of crypts and goblet cells, together with severely diminished levels of mucin (Figure 9C, compare corresponding upper and lower panels). Co-supplementation with DSS/SCN^-^ failed to alter goblet cell damage as colon sections revealed similar levels of pathology between the two DSS groups irrespective of whether SCN^-^ was present or absent (Figure 9C, compare lower panels). Quantification of mucin production normalised to colon length demonstrated a significant reduction in mucin levels for DSS-challenged mice relative to the controls (*p* < 0.05). However, there was no difference in mucin loss for colons from the DSS/Cl^-^ and DSS/SCN^-^ groups (Figure 9D).

### 3.7 Assessing cell apoptosis in the inflamed colon

Colon cell viability, as a biomarker of colitis severity and inflammation, was investigated via a TUNEL assay. Immunohistochemical tagging of DNA fragmentation enables the visualisation of non-viable cells, which were noticeably more prominent across the DSS treatment groups compared to the controls (Figure 10A). Quantification of TUNEL^+^ fluorescent cells in the transverse and descending colon showed markedly higher loss of cell viability in DSS-treated mice than the corresponding controls (*p* < 0.05), but levels of viability did not differ between colons from the DSS/Cl^-^ and DSS/SCN^-^ groups (Figure B).

**Figure 10.**
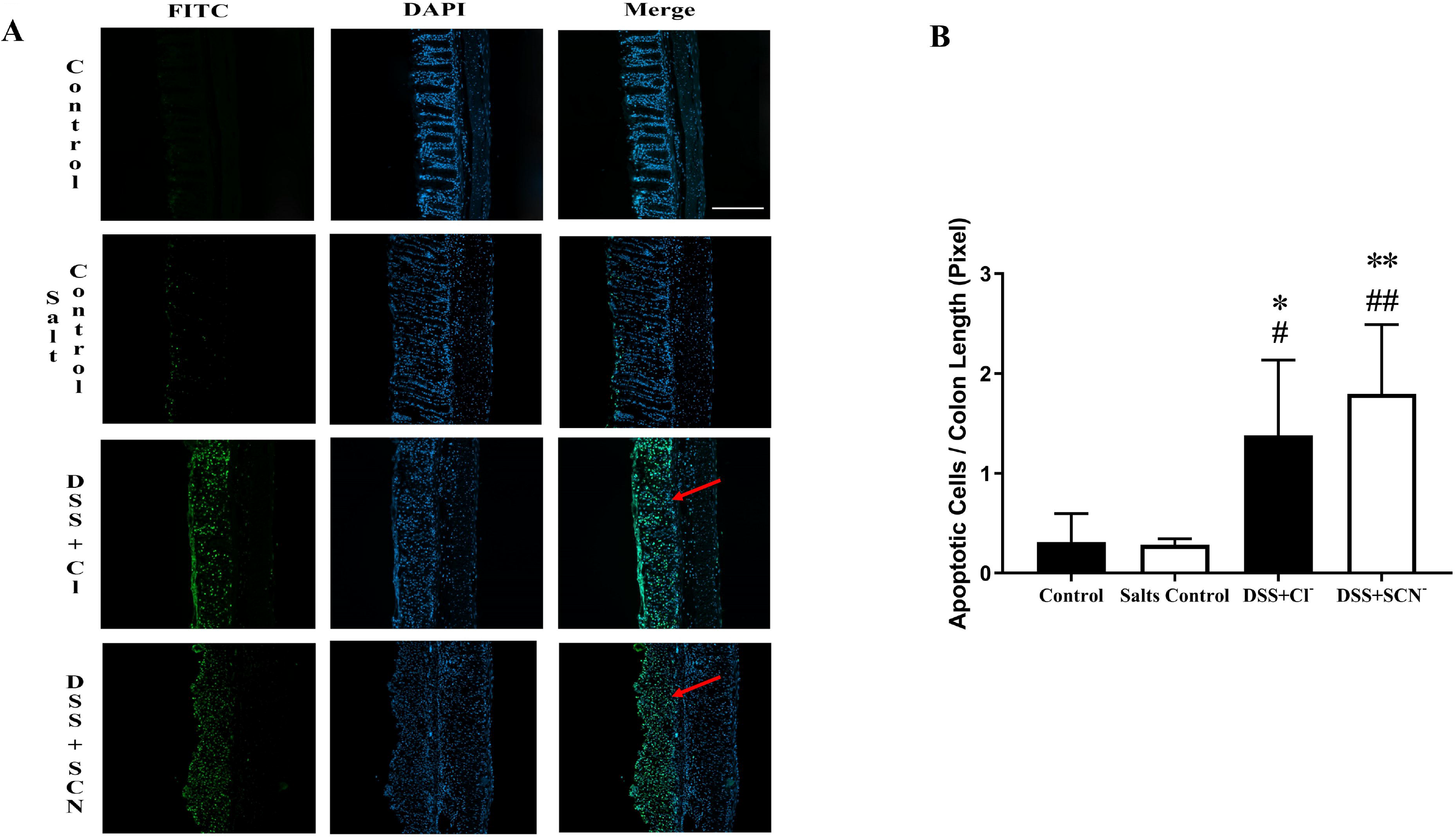
Immunofluorescence labelling and quantification of apoptotic cells in descending colon sections. DNA fragmentations on colon sections are labelled by a fluorometric TUNEL (TdT-mediated dUTP Nick-End Labeling) assay. Panels show (A) representative TUNEL images showing staining of DNA fragmentation (FITC, green, red arrow) and nuclei (DAPI, blue) together with merged/co-registered images. Slides were imaged using the Zeiss AxioScan at 10x magnification. Scale Bar = 100 μm. (B) Quantification of apoptotic (TUNEL^+^) cells adjusted to corresponding pixel length using Metamorph^®^ V 7.1 software. Data represents mean ± SD, n=6 per group for control and DSS, n=5 per group for salts control and DSS + SCN^-^. * Compared to control; \**p* < 0.05, \*\**p* <0.005. ^#^ Compared to salts control; ^#^*p* <0.05, ^##^*p* < 0.005

## 4. Discussion

It is increasingly recognised that oxidative stress and oxidative damage are factors that are linked to perpetuating the cycle of chronic inflammation during active UC [1, 3, 44–46]. Enzymic activity of leukocyte-derived MPO is believed to be the primary source of ROS production in the inflamed colon [44, 47]. Consistent with the study design, dietary SCN^-^ supplementation increased fecal and serum SCN^-^ levels to closely match levels in human smokers [14]. However, SCN^-^ supplementation did not attenuate the course of experimental murine colitis, despite evidence of a trending decrease in MPO 3-Cl-Tyrosine in the colons, perhaps in favor for HOSCN production. Mice activity levels, colon length and colonic histopathological/histoarchitectural were not statistically different between DSS/Cl^-^ and DSS/SCN^-^ groups, although DSS/SCN^-^ mice showed increased lost of bodyweight at Day 9 of DSS challenge, compared to DSS/Cl^-^ mice. No differences were observed in mucin assessment and cell viability between both DSS-treated groups, suggesting no change in the rate of goblet cell dropout or global cell death. Interestingly, mice supplemented with DSS/SCN^-^ showed marked upregulation of Nrf2 and GCLC, indicating that thiol synthesis was enhanced in this group of mice and this may provide an increase in antioxidant status with the colon during DSS-insult. Overall, increasing SCN^-^ in the gut and circulation to match levels in human smokers proved to yield minimal protection against active experimental colitis.

The protective effects of HOSCN is a contentious area of research, with numerous mixed reviews in different disease models. For example, inhaled SCN^-^ supplied via nebuliser to mice with lung infection showed a cytoprotective effect against HOCl and MPO-mediated cell damage, reduced level of infection and associated morbidity [48]. Furthermore, SCN^-^ supplemented in drinking water also decreases atherosclerotic plaque size in human MPO transgenic mice [8]. In humans, elevated plasma SCN^-^ concentrations were associated with improved long-term survival after the first myocardial infarction in a study involving 176 human subjects with current smokers showing much higher long-term survival rate than non-smokers and ex-smokers with low SCN^-^ [18]. On the other hand, there have been reports on the negative effects of SCN^-^ in studies using *in vitro* models, including catalysing MPO-initiated lipid oxidation in different lipoprotein classes [41,42], causing endothelial nitric oxide synthase dysfunction [43] [49]. It is thought that the Zn(2+)-thiol cluster of eNOS are specifically targeted for oxidation by HOSCN, resulting in enzyme dysfunction and reduced formation of NO^•^. Studies in humans have shown excess SCN^-^ caused excessive alteration in plasma thiol levels [18] [14].

The fold increase in SCN^-^ levels in plasma and fecal samples are comparable to those reported for human smokers and other SCN^-^ supplementation studies [8, 16, 50]. Although the study of Morgan and colleagues[8] showed higher SCN^-^ levels in supplemented mice than herein, we anticipate this is due to a longer period of SCN^-^ supplementation (12 weeks as opposed to 9 days). Despite this, we have shown in an *ex vivo* model that ~60% of MPO chlorinating activity was inhibited with 0.1 mM SCN – a similar concentration found in the serum of SCN^-^ treated mice in this study. Chandler and colleagues reported that a HOCl production was completely abrogated at 400 μM SCN^-^ in an *in vitro* model [51]. However, we have shown that the IC_50_ value of NaSCN for inhibition of MPO-mediated luminol oxidation was 0.034 mM, markedly below the levels of SCN detected in serum of SCN^-^-treated mice. If SCN was protective in the murine model of DSS-induced colitis, we would anticipate some attenuation of disease indices with a SCN^-^ serum concentration of ~0.07mM – almost double the IC_50_ value we reported.

Taken together, these data support the delivery of 10 mM SCN^-^ via drinking water was sufficient to cause systemic elevation of endogenous SCN^-^ levels. Quantification of SCN^-^ levels in homogenised colon sample were unable to be performed due to the interferences with ion-exchange chromatography and the investigation of other MPO substrates such as Cl^-^ and Br^-^ were not conducted due to limited availability of plasma samples.

Active smokers, who have been repeatedly shown to have significantly elevated blood SCN^-^ levels, exhibit a protective effect against ulcerative colitis [21–23, 55, 56]. Active smoking reduces the risk for UC by 1.7-fold and patients showed reduced flares, less need for steroids and lower colectomy rate [57]. In addition, many case-controlled studies have demonstrated that ex-smokers exhibit greater risk of UC than non-smokers and current smokers, and this risk is reduced if patients recommence smoking [22, 58, 59]. In a study of 138 UC patients, 70% developed colitis after cessation of smoking for the first year and 52% in the first three years [60].

Hydrogen cyanide (HCN) is found in the mainstream smoke of all cigarettes and upon inhalation, ~ 80% of HCN reacts with a sulfur donor and a sulfur transferase enzyme such as rhodanese to detoxify into thiocyanate ion (SCN^-^) [61]. Multiple large-scale health studies have shown plasma from smokers contained significantly higher endogenous levels of SCN^-^ than that of non-smokers. In non-smokers, plasma SCN^-^ concentration is usually between 10-100 μM, but in smokers, plasma SCN^-^ values have been reported to exceed 300 μM (usually at 180 ± 55) [8, 14, 24, 62]. Thus, smoking-induced plasma SCN^-^ elevation could divert the pattern of oxidative damage in colonic tissue by promoting production of less potent HOSCN over HOCl and may play a beneficial role in the pathophysiology of UC.

The detection of identical levels of calprotectin indicates that SCN^-^ supplementation does not directly modulate colonic MPO levels, parallel to other studies [14, 18]. Calprotectin is consistently expressed in activated neutrophils and macrophages, which are also the primary source of endogenous MPO [4, 63]. Moreover, calprotectin expression in the colon is used as a clinical criterion for measuring the severity of IBD [64]. Thus, detection of calprotectin provides greater depth of knowledge on disease severity in mice than by monitoring MPO alone.

The marked increase in SCN^-^ concentrations in the supplemented mice would be expected to modify the ratio of oxidants produced by MPO, with decreased level of HOCl-related damage and increased HOSCN-related thiol oxidation. In this study, a trending decrease of colonic 3-Cl-Tyr levels, the biomarker of HOCl-specific oxidative damage, reported in SCN^-^ supplemented mice indicate the lack of HOCl and HOCl-mediated oxidation of tyrosine molecules. Thus, SCN^-^ appeared to divert the production of HOCl by MPO to the formation of other oxidants, with HOSCN being the most likely biological outcome. In support of these findings, previously published studies on human plasma samples also showed that high plasma SCN^-^ can increase HOSCN-related thiol oxidation and decrease levels of 3-Cl-Tyr [32]. Surprisingly, thiol oxidation as determined by plasma thiol content of low-molecular weight thiol (glutathione) and protein-bound thiol (PTP) did not reveal a significant difference between DSS-treated mice and their counterparts co-supplemented with NaSCN. This could be due to the localised differences in thiol oxidation in the colon which may not necessarily be reflected by systemic markers, as others have purported with this acute DSS model [8].

Furthermore, compensatory repair mechanisms during DSS–induced colitis may increase thiol turnover, complicating the interpretation of data. Thus, the upregulation of Nrf2 noted here is evidence for an upregulated endogenous antioxidant response [33, 65]. Notably, Nrf2 upregulates many antioxidant genes, including GCLC, the rate-limiting catalysing subunit for GSH formation [41, 43, 66]. Our data demonstrate for the first time that DSS insult stimulates a parallel increase in Nrf2 and GCLC expression in the colon. Moreover, upregulation of the GCLC machinery was noticeably more prominent in SCN^-^ supplemented DSS mice than in the colons of DSS/Cl-supplemented counterparts. This suggests that GSH synthesis was enhanced in SCN^-^ supplemented animals, possibly in response to increased thiol damage by HOSCN, compared to HOCl.

Activation of the transcription factor Nrf2 also stimulates production of a variety of other downstream antioxidants other than GSH in an inflammatory environment such as ulcerative colitis. One example is hemeoxygenase-1 (HO-1) [67, 68]. A possible explanation for lower thiol synthesis in DSS/Cl^-^ mice under similar levels of Nrf2 expression could be that more *non-thiol* antioxidants were oxidised in DSS/Cl^-^ mice. Thus, juxtaposed to the elevated Nrf2 expression in DSS/SCN^-^ mice as a response to GSH depletion, Nrf2 is upregulated in DSS/Cl^-^ mice to stimulate the production of non-thiol antioxidants such as HO-1. Future investigations should be conducted to examine and compare the levels of non-thiol endogenous antioxidants regulated under Nrf2 transcriptional regulation in the two DSS treatment groups as a possible mechanism that alleviates thiol oxidation in the colon.

Ultimately, the diversion of MPO-mediated HOCl formation in the colon to HOSCN did not exert protection in the DSS-induced colitis model herein. This main outcome indicates that SCN^-^ or the production of HOSCN may not play a role in explaining smoking-related protection against ulcerative colitis and point to other components of inhaled smoke as providing protection from colitis pathogenesis. One possibility is nicotine, which is known to exert anti-inflammatory and immunosuppressive effects on the colon and thereby, inhibit inflammation. For example, some studies suggest that nicotine can reduce the production of proinflammatory IL-8 cytokine and increase the levels of anti-inflammatory IL-10 cytokine [69, 70]. Other labs have suggested that nicotine may also act on the neuronal acetylcholine receptor subunit a7 to reduce production of the proinflammatory cytokine tumour necrosis factor [70, 71].

## 5. Conclusion

Collectively, evidence from this study suggests that SCN^-^ supplementation results in increased HOSCN and decreased HOCl production, which diverts the pattern of oxidative damage within the colon as shown by decreased levels of HOCl biomarkers and increased thiol biosynthesis. However, this diversion in the oxidative damage pattern did not result in any clinical and histological difference or amelioration of experimental UC. Thus, the beneficial effect of cigarette smoking to UC protection may not be solely ascribed to preferential HOSCN formation as a result of high SCN^-^ levels but rather other chemical constituents contained within cigarettes.

## Abbreviations

CD: Crohn’s Disease
3-Cl-Tyr: 3-chlorotyrosine
DSS: Dextran Sodium Sulphate
GAPDH: Glyceraldehyde 3-phosphate dehydrogenase
GCLC: Glutamate-Cysteine Ligase Catalytic Subunit
GI: Gastrointestinal
GSH: Glutathione
GSSG: Glutathione Disulfide
HOBr: Hypobromous Acid
HOCl: Hypochlorous Acid
HCN^-^: Hydrogen; Cyanide
HOSCN: Hypothiocyanous Acid
IBD: Inflammatory Bowel Disease
MPO: Myeloperoxidase
Nrf2: Nuclear factor erythroid 2-related Factor 2
PMNs: Polymorphonuclear Neutrophils
PTP: Protein Tyrosine Phosphatases
ROS: Reactive Oxygen Species
SCN^-^: Thiocyanate
UC: Ulcerative Colitis

## Acknowledgements

We thank the Bosch Institute for providing shared facilities for Mass Spectrometry to conduct part of the research herein.

The authors acknowledge funding from the Australian Research Council (grant *DP130103711*) and the NHMRC Project grant (*APP1125392*).

